# Functionalized extracellular matrix scaffolds loaded with endothelial progenitor cells promote neovascularization and diabetic wound healing

**DOI:** 10.1101/2021.02.02.429318

**Authors:** Siqi He, Tanaya Walimbe, Hongyuan Chen, Kewa Gao, Priyadarsini Kumar, Yifan Wei, Dake Hao, Ruiwu Liu, Diana L Farmer, Kit S.Lam, Jianda Zhou, Alyssa Panitch, Aijun Wang

## Abstract

Diabetic ischemic wound treatment remains a critical clinical challenge. Strategies that enhance angiogenesis and improve ischemic pathology may promote the healing of poor wounds, particularly diabetic wounds in highly ischemic condition. We previously identified a cyclic peptide LXW7 that specifically binds to integrin αvβ3 on endothelial progenitor cells (EPCs) and endothelial cells (ECs), activates VEGF receptors, and promotes EC growth and maturation. In this study, we designed and synthesized a pro-angiogenic molecule LXW7-DS-SILY by conjugating LXW7 to a collagen-binding proteoglycan mimetic DS-SILY and further employed this novel bifunctional ligand to functionalize extracellular matrix (ECM) scaffolds, promote neovascularization and accelerate ischemic wound healing. We established a Zucker Diabetic Fatty (ZDF) rat ischemic skin flap model and found the wounds treated by LXW7-DS-SILY-functionalized ECM scaffolds, with or without EPCs, significantly improved wound healing, enhanced neovascularization and modulated collagen fibrillogenesis. These functionalized ECM scaffolds also significantly promoted EPC attachment, growth and survival in the ischemic environment. Altogether, this study provides a promising novel treatment to accelerate diabetic ischemic wound healing, thereby reducing limb amputation and mortality of diabetic patients.

## 1 Introduction

In 2019, the International Diabetes Federation (IDF) estimated a global population of 415 million people with diabetes mellitus, and that the number of people affected will increase to 700 million by 2045 [1]. IDF also estimated that 25% of this population will develop diabetic foot ulcers (DFU) in their lifetime [2], with over 65% of DFUs having an ischemic pathology prone to serious infections, leading to diabetes-related lower extremity amputations [3]. Moreover, ulcers and other foot complications contribute greatly to diabetes-related hospitalizations, and thus represent a significant healthcare burden. Accelerating the wound healing process in ischemic diabetic foot ulcers would reduce the need for lower extremity amputation, improve survival of patients suffering from DFUs, and decrease diabetes-related hospitalizations.

The current standard of care for DFU treatment includes debridement of the wound, antibiotic management of any infection, ulcer off-loading, and revascularization surgeries, if indicated. The current bioengineered scaffold therapies using commercially available scaffolds such as Dermagraft (human fibroblast-derived dermal substitute) and Regranex (recombinant PDGF, also known as becaplermin gel), or tissue engineered skin, but all have disadvantages due to multiple repeat applications for efficacy, secondary surgeries, delayed healing, fibrosis [4]. And none of them target re-vascularization specifically. To overcome these difficulties, stem cell-based therapy to improve revascularization of the tissue has emerged as a new approach for chronic wound healing [5]. Neovascularization plays a significant role in wound healing during all stages of the tissue repair process; therefore, a therapy that improves the vascularization of the damaged and regenerating tissue will hasten the healing process, leading to better outcomes [6].

Endothelial progenitor cells (EPCs) are derived from bone marrow and have the potential to differentiate into mature endothelial cells (ECs), leading to tissue vascularization [7]. Velazquez et al. [8] demonstrated that EPCs are present in much higher numbers in non-ischemic wounds compared to ischemic wounds generated in the same animal. This effect is enhanced in diabetic ischemic wounds, where the lack of access to peripheral blood dampens EPC migration to the wound and leads to impaired healing due to poor oxygenation and nutrient and waste transport [9]. This lack of access and impaired EPC recruitment suggest an unmet need for exogenous EPC delivery to ischemic sites. Localized EPC delivery to the ischemic wound site in biological scaffolds that support EPC engraftment, angiogenesis and wound healing will have a profound beneficial effect on healing in patients with ischemic wounds.

Porcine small intestinal submucosa (SIS) is an FDA-approved, commercially available, collagen-based natural extracellular matrix (ECM) scaffold that has been widely used clinically for various tissue repairs [10]. SIS serves as a suitable provisional matrix for cell delivery and tissue regeneration because of its good biocompatibility and porous structure [11,12]. SIS scaffolds can also be functionalized in many ways and surface modification has been shown to enormously improve its application in tissue engineering [13]. DS-SILY, is a glycan-based therapy designed to mimic decorin, and is composed of collagen-binding peptides (SILY) conjugated to a dermatan sulfate (DS) backbone [14]. Decorin is a small leucine-rich proteoglycan found abundantly in skin, and is involved in maintaining regularity in collagen fibrillogenesis and preventing hypertrophic scars [15–17]. Similar to the functions of native decorin, a key feature of the DS-SILY molecule is that it binds to collagen matrices, thereby protecting the matrix from extensive and rapid proteolytic breakdown [14]. A further drawback of collagen-rich scaffolds is that they lack sufficient innate biological signals to support sufficient recruitment of EPCs and revascularization of ischemic wounds *in vivo*. We previously identified LXW7, a cyclic peptide ligand that binds the αvβ3 integrin receptor on ECs and EPCs, and stimulates cell proliferation and phosphorylation of the VEGF2 receptor [18]. Webber et al. [19] reported that grafting peptides which activate the VEGF receptor to a scaffold results in increased microcirculation and decreased hind-limb ischemia in a mouse model, highlighting the importance of VEGF receptor activation. Hence, the LXW7 peptide ligand could potentially serve to recruit and retain endogenous EPCs by supporting the binding of ECs and EPCs and activating the VEGFR2 receptor through αvβ3 engagement. In order to assemble a stable and functionalized skin tissue engineering scaffold that can provide a temporary residence for maintaining the angiogenic function of EPCs and support cell survival under ischemic condition, we designed a novel molecule, LXW7-DS-SILY, which is composed of both SILY and LXW7 peptides conjugated to dermatin sulfate. LXW7-DS-SILY binds to the collagen present in the SIS scaffold and also binds to EPCs. Here we demonstrate that when LXW7-DS-SILY is added to SIS the new collagen-rich ECM scaffold increased EPC retention at the ischemic site, enhanced angiogenic function, and ultimately led to wound healing (Figure 1).

**Figure 1.**
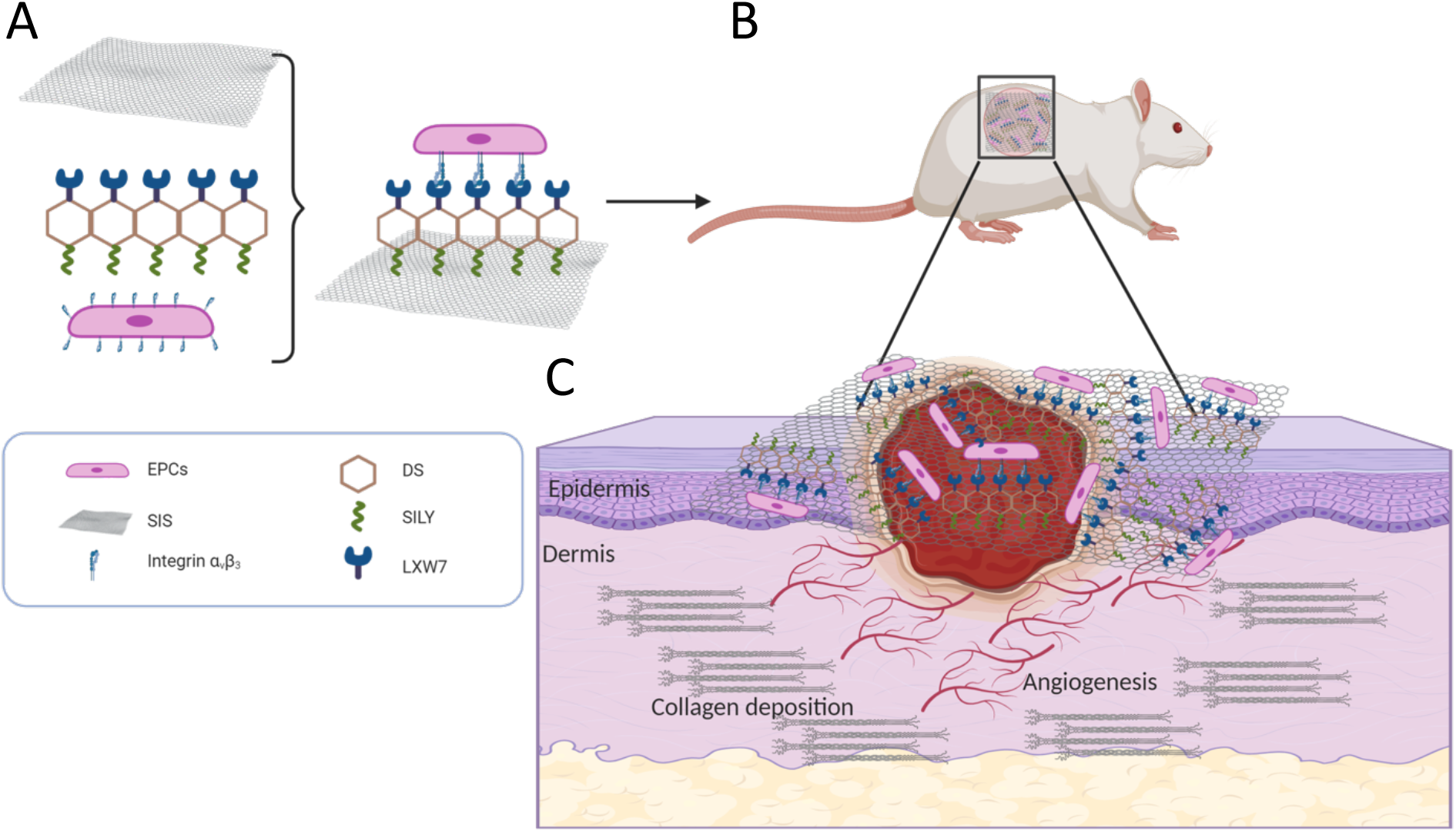
Schematics for the fabrication of ligand modified scaffolds. Multiple functional components combined to generate a functionalized scaffold (**A**) and its topical application to the site of the diabetic ischemic wound of the rat (**B**) leading to improved vascularization and hastening of the healing of the damaged tissue (**C**).

## 2 Materials and methods

### 2.1. Cell characterization and expansion

Bone marrow-derived endothelial progenitor cells (EPCs) of Zucker Diabetic Fatty (ZDF) rats were purchased from Cell Biologics, Inc. (RD-6031). EPCs were characterized by DiI-Ac-LDL staining, immunostaining of CD31 and VE-Cadherin, and tube formation assay. EPCs were cultured in complete rat endothelial cell medium (M1266, Cell Biologics). EPCs were used between passages 5 and 8 for all the experiments described in this study.

#### 2.1.1 Lentiviral vector transduction

The lentiviral construct was generated at the University of California Davis Institute for Regenerative Cures (IRC) Vector Core. EPCs were transduced with pCCLc-MNDU3-LUC-PGK-Tomato-WPRE lentiviral vector in transduction media consisting of rat basal endothelial cell medium, 5% FBS, and 8 µg/ml protamine sulfate (MP Biomedicals) at a multiplicity of infection (MOI) of 10 for 6 h. The transduction medium was then replaced with complete rat endothelial cell medium and the cells were cultured for additional 72 h.

#### 2.1.2 Acetylated low-density lipoprotein uptake assay

EPCs were cultured in serum-free medium for 12 h and then incubated with 10 µg/ml Dil-Ac-LDL (Alfa Aesar) for 5 h at 37 °C, 5% CO2. Cells were then washed three times with Dulbecco’s Phosphate buffered saline (DPBS, HyClone) and fixed with 10% formalin (ThermoFisher Scientific) for 15 min. The nuclei were stained with DAPI (ThermoFisher Scientific) for 5 min. Cells were washed 3 times and imaged with a Zeiss Observer Z1 microscope.

#### 2.1.3 Tube formation assay

A 96-well plate was coated with 50 µl Matrigel (354234, Corning) per well and incubated at 37 °C, 5% CO2 to gel for 60 min. 1 × 10^4^ EPCs were seeded onto the Matrigel-coated wells and incubated at 37 °C for 16 h. Phase contrast images were taken using Zeiss Observer Z1 microscope.

#### 2.1.4 Immunofluorescent staining of rat EPCs

Cells were fixed in 10% formalin for 10 min and non-specific binding sites were blocked with 1% bovine serum albumin (BSA, bioWORLD) in 1X DPBS for 1 h at room temperature (RT). The cells were then stained with either CD31 (Abcam, ab119339, 1:200) or VE-Cadherin (ThermoFisher Scientific, 36-1900, 1:200) antibodies in 1% BSA in DPBS and incubated overnight at 4 °C. Subsequently the cells were then incubated with Alexa488 or Alexa647 conjugated secondary antibodies (ThermoFisher Scientific, 1:500) for 1 h at RT. The cell nuclei were stained with DAPI for 5 min and imaged using Zeiss Observer Z1 microscope.

### 2.2 EPC binding assay on LXW7-beads

For the on bead cell binding assay, 1 × 10^5^ EPCs were added to an ultralow attachment 24-well plate followed by addition of LXW7 resin beads. The plate was placed on shaker set at 40 rpm and incubated for 15 min at 37 °C, 5% CO2. Phase contrast images were taken using Zeiss Observer Z1 microscope.

### 2.3 Flow cytometry analysis of ligand–cell binding affinity

To quantitatively determinate ligand-cell binding affinity, flow cytometry analysis of LXW7-biotin (ligand) or D-biotin (negative control) bound to EPCs was performed, as described in our previous study [18]. Briefly, 3 × 10^5^ EPCs were incubated with 1 µM LXW7-biotin in a binding buffer (1X HEPES (Gibco) containing 10% FBS (HyClone) and 1 mM Mn^2+^) on ice for 30 min. The samples were washed three times with wash buffer (DPBS containing 1% FBS) and incubated with 2 µg/ml of streptavidin-PE-Cy7 conjugate (Life Technologies) on ice for 30 min and then washed with wash buffer. To test the expression of the αvβ3 integrin on EPCs, samples were stained with 20 µg/ml mouse anti-rat αvβ3 integrin antibody (Millipore Sigma) on ice for 30 min, washed three times with wash buffer, and incubated with donkey anti-mouse Alexa647 conjugate (1:500, ThermoFisher Scientific) in DPBS on ice for 30 min and then washed with DPBS. To confirm that LXW7 binds to EPCs mainly via αvβ3 integrin, we performed a binding/blocking experiment using a monoclonal anti-αvβ3 integrin antibody. To block αvβ3 integrin, cells were first incubated with 20 µg/ml mouse anti-rat αvβ3 integrin antibody on ice for 30 min, washed three times with wash buffer, and then incubated with LXW7-biotin (1 µM) for another 30 min. The samples were washed three times with wash buffer and incubated with 2 µg/ml Streptavidin-PE-Cy7 conjugate (Life Technologies) in DPBS on ice for 30 min and then washed with DPBS. Streptavidin-PE-Cy7 only was used as the negative control. Attune NxT Flow Cytometer (ThermoFisher Scientific) was used to perform flow cytometry, and data was analyzed using FlowJo software (FlowJo LLC). The blocking efficiency of LXW7 to ZDF-EPCs was calculated according to the following formula: blocking efficiency of LXW7 to ZDF-EPCs = (B0–Bb)/B0*100, where B0 is designated as the initial ZDF-EPC-LXW7 binding percentage and Bb is designated as the ZDF-EPC-LXW7 binding percentage after ZDF-EPCs were blocked with anti-αvβ3 integrin antibody.

### 2.4 Synthesis and characterization of peptide-hydrazides

Hydrazide-modified peptides RRANAALKAGELYKSILYGSG-hydrazide (SILY-hydrazide) and cGRGDdvc(AEEA)2WG-hydrazide (LXW7-hydrazide, wherein AEEA is short PEG linker) were synthesized using standard Fmoc solid-phase peptide synthesis according to a previously published protocol [20, 21]. Briefly, Cl-TCP(Cl) ProTide Resin (loading 0.4-0.6 mmol/g, CEM Corporation) was rinsed 3 times with dichloromethane (DCM, Fisher Scientific) and N, N-Dimethylformamide (DMF, Fisher Scientific) and swollen in 50% DCM/DMF for 1 h. Swollen resin was then reacted twice with 10% hydrazine hydrate (Sigma) in DMF and 0.057 M N, N-Diisopropylethylamine (DIPEA, Fisher Scientific) for 2 h at RT each. 10% methanol (Fisher Scientific) in DMF was used to cap any unreacted chloride groups, and the resin was washed 3 times with DMF and 3 times with DCM. The resin was then reacted with 4 equivalents of the first Fmoc-amino acid with 4 equivalents of OxymaPure, N,N′-Diisopropylcarbodiimide (DIC, Fisher Scientific) and 10 equivalents of DIPEA in DMF for 4 h, followed by washing three times with DMF and three times with DCM. Subsequent amino acids were coupled for 10 min each at 50 °C on a Liberty Blue microwave peptide synthesizer (CEM Corporation) using 5 equivalents of Fmoc-amino acids, DIC, and OxymaPure with 0.1M DIPEA. 20% piperidine in DMF was used for deprotection. Peptides were cleaved off beads using 88% trifluoroacetic acid (TFA, Fisher Scientific), 5% phenol (Sigma), 5% H2O, and 2% Triisopropylsilane (TIPS, Sigma) for 3 h. Crude peptides were precipitated in cold diethylether (Acro organics) and allowed to dry before dissolving in 5% acetonitrile/H2O for purification. Before purifying, LXW7 was cyclized by oxidizing cysteine residues using ClearOx resin (Peptides International) according to manufacturer’s protocol. Peptides were purified through a C18 prep column against an acetonitrile gradient on an AKTApure 25 FPLC (GE Healthcare) and confirmed by MALDI-TOF-MS (Bruker). For some experiments, peptides were purchased from InnoPep.

### 2.5 Synthesis and characterization of molecule variants

DS-SILY or LXW7-DS-SILY were prepared by conjugating SILY-hydrazide and LXW7-hydrazide to a dermatan sulfate (DS) backbone using carbodiimide chemistry. DS (average molecular weight 41,816 Da, Celsus Laboratories) was reacted with peptide-hydrazides using 1-ethyl-3-[3-dimethylaminopropyl] carbodiimide hydrochloride (EDC, ThermoFisher Scientific) in 0.1 M MES [2-(N-morpholino)ethanesulfonic acid] buffer with 8 M urea (Sigma) and 0.6% NaCl (Sigma) titrated to pH 4.5. First, SILY-hydrazide was conjugated to the DS for 4 h using 0.01 mM EDC. The reaction was stopped by titrating the pH to 8. The product was purified using tangential flow filtration (Spectrum labs) with a 10kDa column and then lyophilized. LXW7-hydrazide was then sequentially conjugated to the DS-SILY construct in a similar manner using 0.1 mM EDC for 24 h before purification. Peptide conjugation was verified by creating standard curves for SILY and LXW7 using concentration dependent 280 nm absorbance of aromatic amino acids and extrapolating absorbances of synthesized molecules using readings taken on a NanoDrop UV-Vis spectrophotometer (Thermo Fisher). The structures of LXW7-DS-SILY and the reaction scheme of making LXW7-DS-SILY was described in a previous study [22].

### 2.6 Preparation of ligand-modified SIS scaffolds with or without EPCs

8-mm diameter punch-outs of SIS-ECM (Cook Biotech) were cut using a sterile biopsy punch and placed into wells of a 48-well plate with the rough side facing up and incubated with 0.2 ml of 10 µM LXW7-DS-SILY, DS-SILY or DPBS for 1 h at 37 °C. Scaffolds were subsequently rinsed with DPBS for three times and then soaked overnight in complete culture media at 37 °C, 5% CO2. 5 × 10^5^ Td-Tomato/luciferin-labeled EPCs were resuspended in 15 µl of complete media per ECM and carefully added onto the surface of LXW7-DS-SILY, DS-SILY ligand modified or unmodified SIS. The plate was placed in a 37 °C, 5% CO2 incubator and incubated for 1 h to allow for cell adherence before addition of 0.3 ml of complete media per well.

### 2.7 Attachment assay and CCK-8 assay

To modify the culture surface with ligands, plates were coated with 200 µl of 20 µg/ml Avidin (Thermo Fisher) and incubated for 1 h at 37 °C. Avidin coated wells were rinsed three times with DPBS and were treated with 200 µl molar equivalents (2 µM) of LXW7-biotin or D-biotin (negative control). The wells were washed three times with DPBS in 1h and blocked with 1% BSA for 1 h. After the wells were rinsed three times with DPBS, 2 × 10^4^ EPCs were added to the wells and incubated for 20 min at 37 °C and 5% CO2. To further explore the effect of LXW7 to the binding of EPCs on an ECM scaffold surface, LXW7-DS-SILY modified SIS and untreated control SIS were placed in an ultralow attachment 48-well plate. The scaffolds were rinsed with DPBS and seeded with EPCs at a density of 1.5 × 10^5^ cells/cm^2^. After 20 min of incubation, unattached cells were washed off with DPBS three times. Images were taken using Zeiss Observer Z1 microscope. Three independent experiments were performed. Cell counter ImageJ plugin was used to determine the cell numbers in three randomly selected fields from each independent experiment.

The effect of LXW7 on the EPC growth was determined by a Cell Counting Kit-8 assay (CCK-8, Dojindo). This assay was used to detect cell viability. The 96-well plates were treated with 1 µM Avidin solution for 1 h at RT then rinsed three times with DPBS. Wells were treated with 1 µM LXW7-biotin (ligand) or 1 µM D-biotin (negative control) for 1 h, washed three times with DPBS and blocked with 1% BSA for 1 h. A total of 1.5 × 10^3^ viable EPCs were seeded per well of the 96-well plate and cultured for 5 days. To test the growth of EPCs on LXW7-DS-SILY modified SIS scaffolds, we prepared the ligand modified scaffolds, as described above in section 2.6. A total of 5 × 10^4^ viable EPCs were seeded per scaffold and cultured for 5 days. The CCK-8 assay was performed according to the manufacturer’s instructions. Absorbance was measured at 450 nm using a SpectraMax i3 plate reader instrument (Molecular Devices LLC).

### 2.8 Cell loading density

The optimal loading density of EPCs on SIS-ECM was determined by the CCK-8 assay. EPCs were seeded on 8-mm SIS-ECM discs at the following densities: 0, 4.2 × 10^4^, 1 × 10^5^, 3 × 10^5^, 5 × 10^5^, and 1 × 10^6^ cells/cm^2^. At 24 h after seeding, absorbance was measured at 450 nm.

### 2.9 Zucker Diabetic Fatty rat ischemic skin wound model

All animal procedures were approved by The University of California, Davis (UCD) Institutional Animal Care and Use Committee (IACUC). The ligand-modified SIS scaffolds loaded with or without EPCs were prepared as described above in methods section 2.6. Male Zucker Diabetic Fatty rats (ZDF fa/fa, Charles River, 12 to 16 weeks old) were used and placed on a special diet to induce Type II diabetes (Purina #5008 from Lab Diet, Inc.). The blood glucose levels of ZDF rats in each group were measured using a digital glucose meter and test strips (Accu-Chek® Sensor, Roche Inc., Mannheim, Germany) 2 or 3 days prior to surgery and at the study endpoint. The rat ischemic skin wound model was carried out as previously described with some modifications [23]. Briefly, animals were placed under isoflurane anesthesia (5% induction) and maintained (1-3%) on a non-rebreathable circuit with charcoal filter scavenging. After shaving and sterilization, two 6-mm diameter full-thickness ischemic wounds were created using a biopsy punch in the center of the designated flap area (Figure S1, 10 cm × 3 cm). Incisions were then made along the marked lines down to the paraspinous muscle to create a bipedicle flap and separate the panniculus carnosus fascia from the paraspinous muscles. A sheet of sterile silicone was placed underneath to induce ischemia. 5-0 non-absorbable sutures (J494G, eSutures) were used to close both incisions by anchoring the silicone sheet to the skin with multiple interrupted stitches along the length of the flap. Lastly, two 6-mm diameter full-thickness non-ischemic wounds were created using a biopsy punch lateral to the bipedicle flap on either side of the flap. The wounds created were then covered with the following SIS-scaffold treatment groups: SIS only, SIS/DS-SILY, SIS/LXW7-DS-SILY, SIS/EPCs, SIS/DS-SILY/EPCs, or SIS/LXW7-DS-SILY/EPCs (prepared as described in section 2.6). The EPCs-loaded SIS scaffolds were placed with cells in direct contact with the site of the wound. Scaffolds were immobilized with 4 stitches using 5-0 silk sutures (VP710X, eSutures). To protect the wound, a Tegaderm transparent dressing (3M Health Care, Germany) was then applied to all wounds. At days 0, 3, 7, 11, and 14 post-treatment, pictures were taken of all wounds, blinded for treatment type. The wound area was calculated by tracing the wound margins from rats and was evaluated as a percent area of the original wound using Image-Pro Plus 6.0 software. The percentage of wound reduction was calculated according to the following formula. Rate of wound closure = (A0-At)/A0*100, where A0 and At are designated as the initial wound area and wound area at the designated time, respectively.

### 2.10 Bioluminescence imaging

Scaffolds seeded with EPCs in the ZDF rats were monitored via In Vivo Imaging Spectrum (IVIS) system (PerkinElmer) at designated time points (1, 3, 7 and 11 days post-treatment). Prior to imaging, the rats were weighed, and luciferase substrate D-luciferin (Gold Biotechnology) was injected subcutaneously into the animals at 150 mg/kg body weight. Five min post-injection, the rats were anesthetized using 2% isoflurane for 5 min and imaged under anesthesia. Images were analyzed using Living Image^®^2.50 (Perkin Elmer). The bioluminescence signal intensity was represented as total photons of the region of interest (ROI) subtracted by the region of no positive signal in the same animal.

### 2.11 Histological analysis

Animals were euthanized on day 14 and their wound tissues were carefully biopsied. Samples were fixed with 4% paraformaldehyde for 24 h, protected by 30% sucrose dehydration for 48 h, and embedded in O.C.T compound (Sakura Finetek USA). Sections (10 µm thick) were cut using a Cryostat (Leica CM3050S) and stained with Hematoxylin and Eosin (H&E) to visualize tissue formation or Masson trichrome staining to observe collagen deposition during the healing period. A total of 24 single images were captured at 10X magnification, merged and analyzed using a Keyence BZ9000 microscope and the BZ II Analyzer software. A total of 5 sections from 5 rats of each group were analyzed.

For immunofluorescence staining, tissue sections were washed with DPBS, blocked with 5% BSA for 1 h in DPBS, and stained with the following primary antibodies by incubating at 4 °C overnight: RECA-1 (1:50, ab9774, mouse, Abcam), α-SMA (1:200, ab5694, rabbit, Abcam), Collagen I (1:200, ab21286, rabbit, Abcam), Collagen III (1:200, ab7778, rabbit, Abcam). Sections were then incubated with their respective secondary antibodies diluted at 1:500 for 1 h. The slides were counterstained with DAPI (1:5,000) for 5 min and mounted with Prolong Diamond Antifade Mountant (Invitrogen). A Nikon A1 laser-scanning confocal microscope was used to acquire confocal images. The number of blood vessels (RECA positive) per field were counted and mean fluorescence intensity were measured by Image J. A total of 6 sections from 3 rats of each group were analyzed.

### 2.12 Statistics

Data are reported as mean ± standard deviation (SD) for bead-binding, block efficiency, cell attachment, CCK-8 assay and cell loading density assay and as mean ± standard error of mean (SEM) for healing rate, bioluminescence images and histological analysis. Statistical analysis of beads binding assay and cell attachment assay was performed using the Student’s t-test. Analyses of bioluminescence images, histological analysis were performed using one-way ANOVA. Healing rate analysis, CCK-8 assay was performed by two-way ANOVA. All statistical analyses were performed using PRISM 7 (GraphPad Software Inc.), and differences were considered significant when p < 0.05.

## 3 Results

### 3.1. Characterization and transduction of ZDF rat bone marrow EPCs

ZDF rat bone marrow EPCs (ZDF-EPCs) displayed the typical morphology of EPC colonies (Figure S2A) and were efficiently transduced with lentiviral vector expressing a fluorescent marker Td-tomato for tracking analysis (Figure S2B). Strong expression of EPC markers (CD31 and VE-cadherin) confirmed their EC characteristics (Figure S2C-D). The tube formation assay results revealed their ability to assemble into tubules in the presence of a basement membrane *in vitro* (Figure S2E). After co-culturing with DiI-Ac-LDL, positive staining for DiI-Ac-LDL was observed in ZDF-EPCs (Figure S2F). These results indicated that ZDF-EPCs have similar phenotype and function as human EPCs.

### 3.2 LXW7 ligand showed excellent binding affinity to ZDF-EPCs via the αvβ3 integrin

In our previous studies, we identified LXW7 as a potent ligand that specifically targets EPCs and ECs normally present in circulation via binding to αvβ3 integrin on cell surfaces [18, 24]. To confirm the binding of LXW7 to the ZDF-EPCs, resin beads displaying LXW7 were incubated with ZDF-EPCs. THP-1 monocytes served as a negative control. After 30 min of incubation, the LXW7 beads showed strong binding to ZDF-EPCs but not to THP-1 monocytes (Figure 2A). Quantification of the number of cells bound on each bead showed that there was a significant increase in the binding of ZDF-EPCs to LXW7 resin beads compared to THP-1 monocytes (Figure 2B). To further confirm that LXW7 binds to ZDF-EPCs mainly via αvβ3 integrin, we performed flow cytometry analysis to detect the expression of αvβ3 integrin and performed a binding/blocking experiment using a monoclonal anti-αvβ3 integrin antibody. The results showed αvβ3 integrin was expressed on the majority of ZDF-EPCs [(75.33±4.25% of overall populations) Figure 2C]. From the binding/blocking experiment, as shown in Figure 2D, we confirmed that LXW7 had high binding efficiency to ZDF-EPCs, and LXW7 binding to ZDF-EPCs was blocked by the anti-αvβ3 integrin antibody (Figure 2D). The average blocking efficiency was 60.8±22.1% (Figure 2D). These data confirmed that the binding of LXW7 to ZDF-EPCs was primarily mediated via αvβ3 integrin.

**Fig 2.**
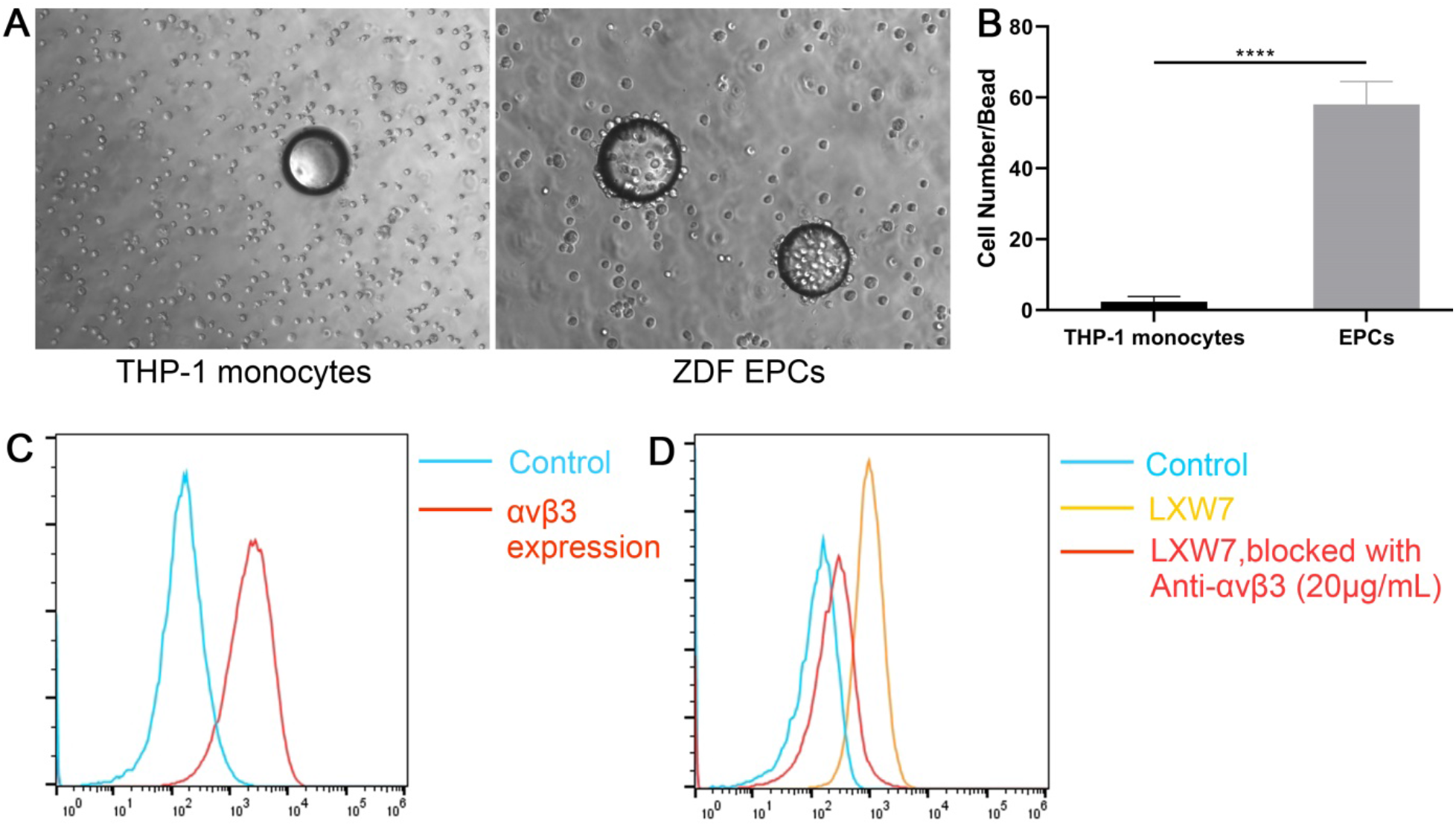
Specificity and binding affinity of LXW7 ligand to ZDF-EPCs and expression of αvβ3 integrin on ZDF-EPCs A. On-bead cell binding assay for testing the binding affinity and specificity of ligands to ZDF-EPCs showing high binding of ZDF-EPCs to LXW7 coated beads (right panel) compared to THP-1 monocytes (left panel). **B**. Quantification of the number of THP-1 monocytes and ZDF-EPCs bound to the LXW7 coated beads. **C**. Expression of αvβ3 integrin on ZDF-EPCs. **D**. Flow cytometry analysis of the binding affinity of LXW7 showing the blocking of its binding by anti-αvβ3 antibody. Scale bar = 100 µm. Data are expressed as mean ± SD. ****p<0.0001, n=3.

### 3.3 LXW7 modified surfaces increased ZDF-EPC attachment and growth

To determine if the LXW7 ligand can support effective ZDF-EPC attachment and growth, we employed both 2D tissue culture surface and 3D ECM-based scaffolds to determine cell-ligand binding ability in different situations. We treated tissue culture polystyrene with LXW7-biotin (ligand) or D-biotin (negative control) and investigated selective attachment of ZDF-EPCs. To further explore the binding ability of LXW7 to ZDF-EPCs on ECM-based scaffolds, we incubated LXW7-DS-SILY with SIS to allow binding of the biofunctional ligand to the SIS. Seeding ZDF-EPCs on both the 2D and 3D surfaces showed that LXW7 and LXW7-DS-SILY can support attachment of ZDF-EPCs both to plate and SIS surfaces (Figure 3A, 3B, 3D, 3E). The number of cells attached was quantified and the results demonstrated that the LXW7-treated surfaces attracted more ZDF-EPCs than the unmodified surfaces (Figure 3C, 3F respectively).

**Figure 3.**
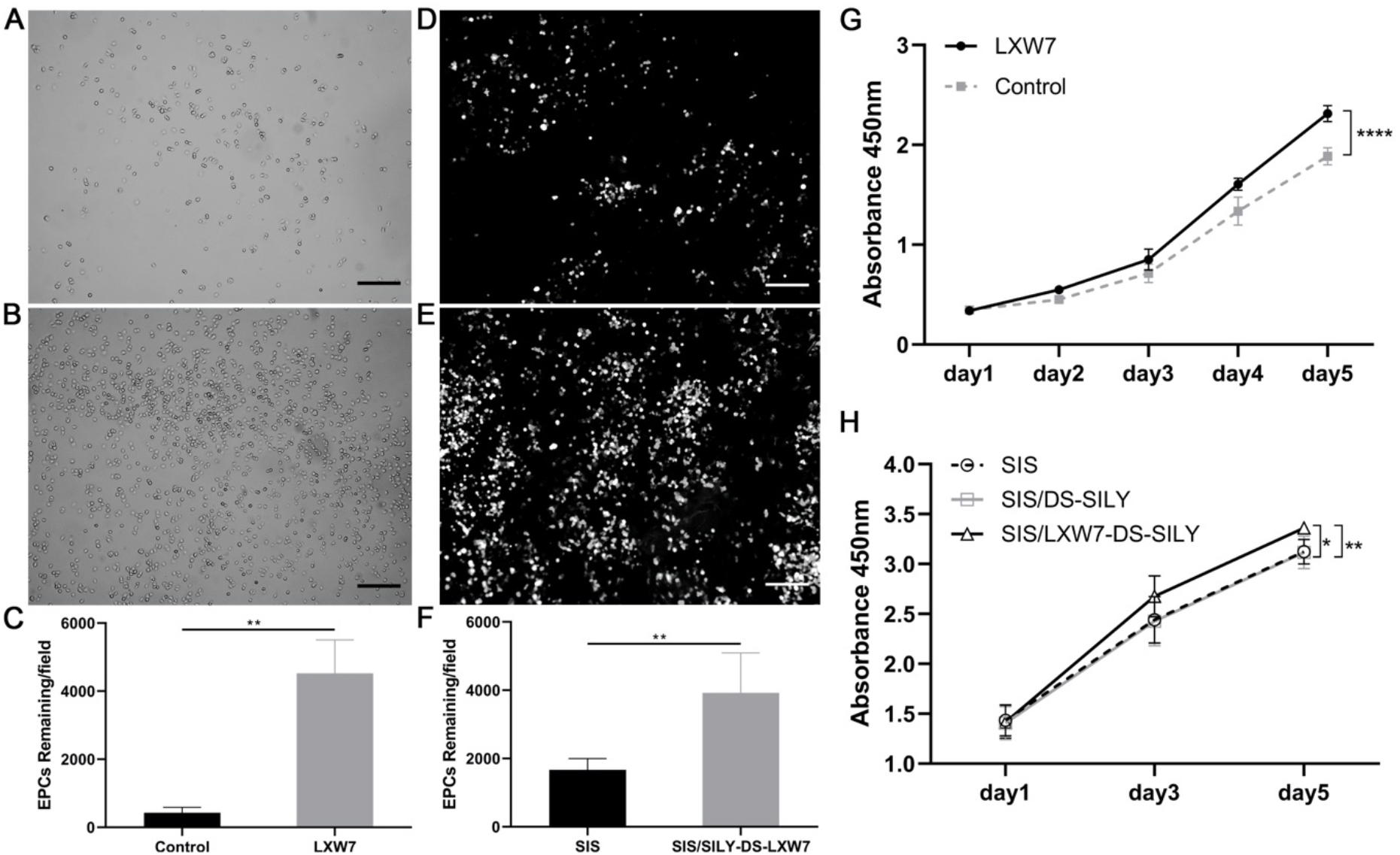
Effect of LXW7 and LXW7-DS-SILY ligands on the attachment, growth and viability of ZDF-EPCs on ligand modified surfaces. **A-B**. Representative images of attached ZDF-EPCs on surfaces treated by D-biotin (A; control), LXW7 after 20 min incubation. (**C**). Quantification and the correlative statistical analysis of remaining cells shown in **A-B. D-E**. Representative images of adhered EPCs on untreated (**D**; control) or LXW7-DS-SILY modified SIS surface (**E**) after 20 min incubation. **F**. Quantification and the correlative statistical analysis of remaining cells shown in **D-E. G**. Growth and viability of ZDF-EPCs on LXW7 treated tissue culture surfaces and D-biotin treated surfaces (control) was assessed by CCK-8 assay. **H**. Growth and viability of ZDF-EPCs on LXW7-DS-SILY or DS-SILY treated SIS scaffold surfaces was assessed by CCK-8 assay. Scale bar = 200 µm. Data are expressed as mean ± SD. ****p<0.0001, **p<0.01, *p<0.05, n=3.

The effect of LXW7 on the growth of ZDF-EPCs was tested by CCK-8 assay at different time points in a course of 5 days. Results showed that the LXW7-biotin treated surface significantly promoted the growth of ZDF-EPCs after 48 h in culture compared with the D-biotin treated surface (control), and this trend was maintained for the entire period of the experiment (Figure 3G). Similar to the trend on tissue culture plates, SIS/LXW7-DS-SILY also promoted the growth of ZDF-EPCs when compared to SIS/DS-SILY and SIS (Figure 3H). These results demonstrate that the LXW7-modified surface and functionalized scaffold support attachment and growth of EPCs.

### 3.4 *In vivo* wound healing studies

A bipedicle cutaneous flap was created on each rat to yield 2 ischemic and 2 non-ischemic wounds per animal (Figure S1A, S1B). Both ischemic and non-ischemic wounds were implanted with ligand-modified or unmodified SIS patches seeded with or without ZDF-EPCs at a seeding density of 5 × 10^5^ cells/cm^2^. This seeding density was based on the result obtained from our CCK-8 assay where a decrease in cell viability was observed beyond a density of 5 × 10^5^ cells/cm^2^ (Figure S3). The average blood glucose level of ZDF rats in each group > 250 mg/dl.

#### 3.4.1 LXW7-DS-SILY modified SIS scaffolds accelerated diabetic non-ischemic wound healing and supported survival of ZDF-EPCs

The diabetic wound healing assay was performed to evaluate the effect of the different scaffolds on non-ischemic wound healing *in vivo*. The wound regions of the diabetic rat were treated with SIS/LXW7-DS-SILY/EPCs, SIS/DS-SILY/EPCs, SIS/EPCs, SIS/LXW7-DS-SILY, SIS/DS-SILY and SIS respectively. Digital photographs were obtained 0, 3, 7, 11, and 14 days after surgery (Figure 4A). In the absence of exogenous ZDF-EPC seeding, the wound area in all groups decreased over time, and the average observed wound closure rate was significantly increased in the SIS/LXW7-DS-SILY group (80.7±0.03%) compared to both the SIS/DS-SILY (56.9±0.01%) and SIS (56.3±0.05%) groups on day 14 (Figure 4B panel a). Among all groups loaded with exogenous ZDF-EPCs, the group treated with SIS/LXW7-DS-SILY/EPCs showed a wound healing rate of 82.5±0.02%, significantly higher than that of SIS/DS-SILY/EPCs (69.7±0.03%) and SIS/EPCs (67.2±0.03%) groups (Figure 4B panel b). However, in the non-ischemic wounds, ZDF-EPCs did not accelerate wound closure when comparing SIS/LXW7-DS-SILY/EPCs with SIS/LXW7-DS-SILY or comparing SIS/EPCs with SIS. When compared to SIS/DS-SILY, SIS/DS-SILY/EPCs accelerated wound healing. (Figure 4B panels a-e).

**Figure 4.**
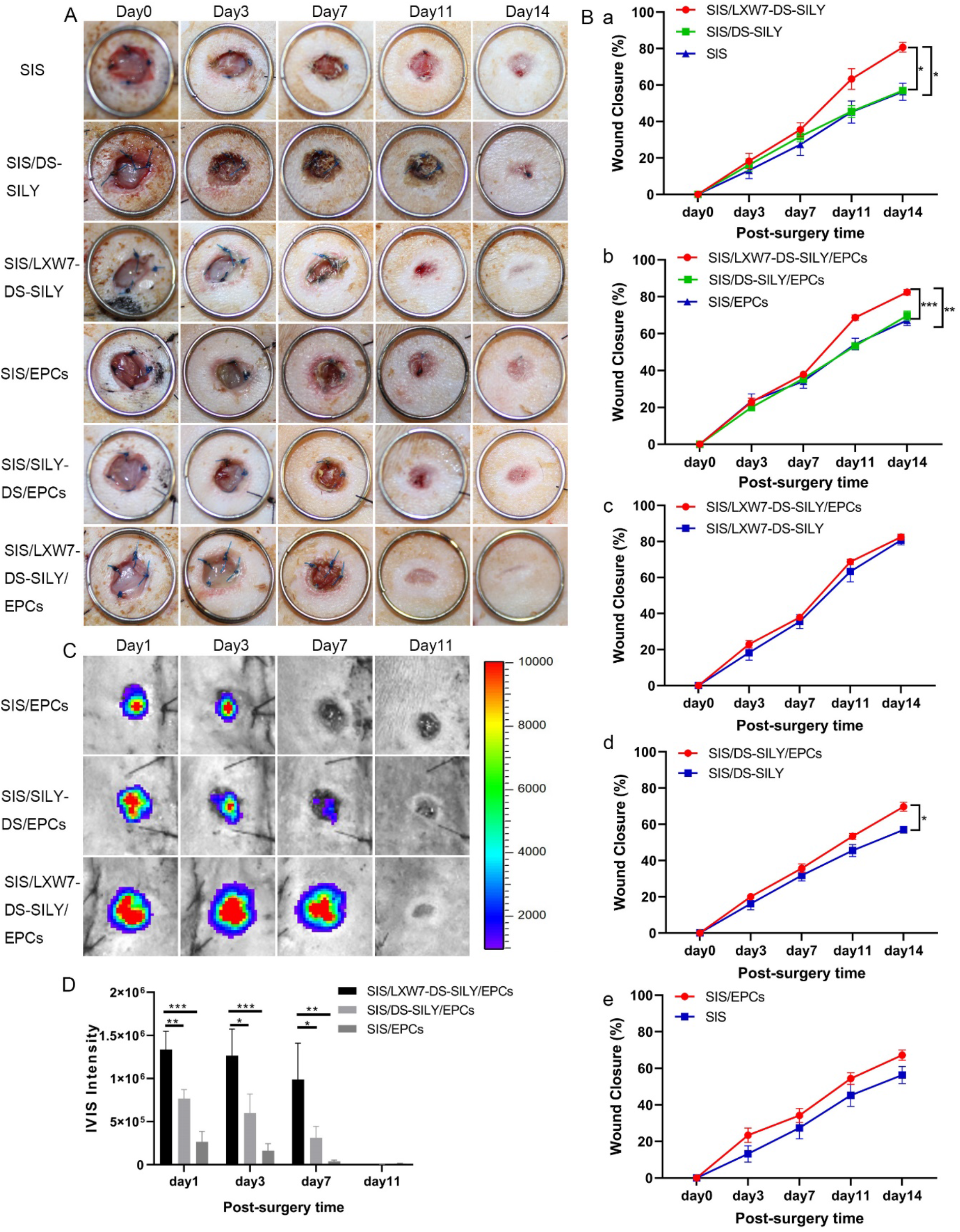
SIS/LXW7-DS-SILY scaffolds accelerate wound closure and support ZDF-EPC survival under non-ischemic condition. **A**. Representative images of healing in wounds treated with different groups at day 0, 3, 7, 11 and 14. **B**. Quantification of the healing rate of different groups. Data are expressed as mean ± SEM. SIS, SIS/LXW7-DS-SILY, SIS/EPCs group: n=8; SIS/DS-SILY: n=7; SIS/DS-SILY/EPCs: n=12; SIS/LXW7-DS-SILY/EPCs: n=13. **C**. Bioluminescence imaging of wounds transplanted with different modified SIS scaffolds loaded with Td-Tomato/luciferin-labeled ZDF-EPCs. **D**. Quantification of the signal intensity of bioluminescence (Most of the SIS scaffolds spontaneously fall off the wounds by day 11). Data are expressed as mean ± SEM. n=5 per group. ***p<0.001, **p<0.01, *p<0.05.

To investigate cell survival of transplanted cells in ZDF rats, Td-Tomato/luciferin-labeled ZDF-EPCs were seeded on different scaffolds and monitored by IVIS. The bioluminescence signal in the SIS/LXW7-DS-SILY/EPCs group showed a significantly higher intensity when compared to SIS/DS-SILY/EPCs and SIS/EPCs groups at day 1, 3 and 7, suggesting that LXW7 improved ZDF-EPC retention in the wound bed at early time points (Figure 4C). The intensity in all the three groups decreased to baseline by day 11 (Figure 4D).

#### 3.4.2 LXW7-DS-SILY modified SIS scaffolds accelerated wound healing and re-epithelialization in non-ischemic wound areas

To further compare the healing quality of wounds treated by different scaffolds, H&E staining was performed on non-ischemic wound tissue samples (Figure 5A). As shown in Figure 5C, the SIS/LXW7-DS-SILY/EPCs group showed the shortest residual wound length among all the groups, followed by SIS/LXW7-DS-SILY, SIS/DS-SILY/EPCs, SIS/EPCs, SIS and SIS/DS-SILY groups at day 14. Also, the SIS/LXW7-DS-SILY/EPCs and SIS/DS-SILY/EPCs groups were fully covered with neoepidermis, while in the other groups, re-epithelialization was not fully completed (Figure 5D). Overall, the SIS/LXW7-DS-SILY/EPCs group showed a higher efficacy of wound healing in comparison with other groups.

**Figure 5.**
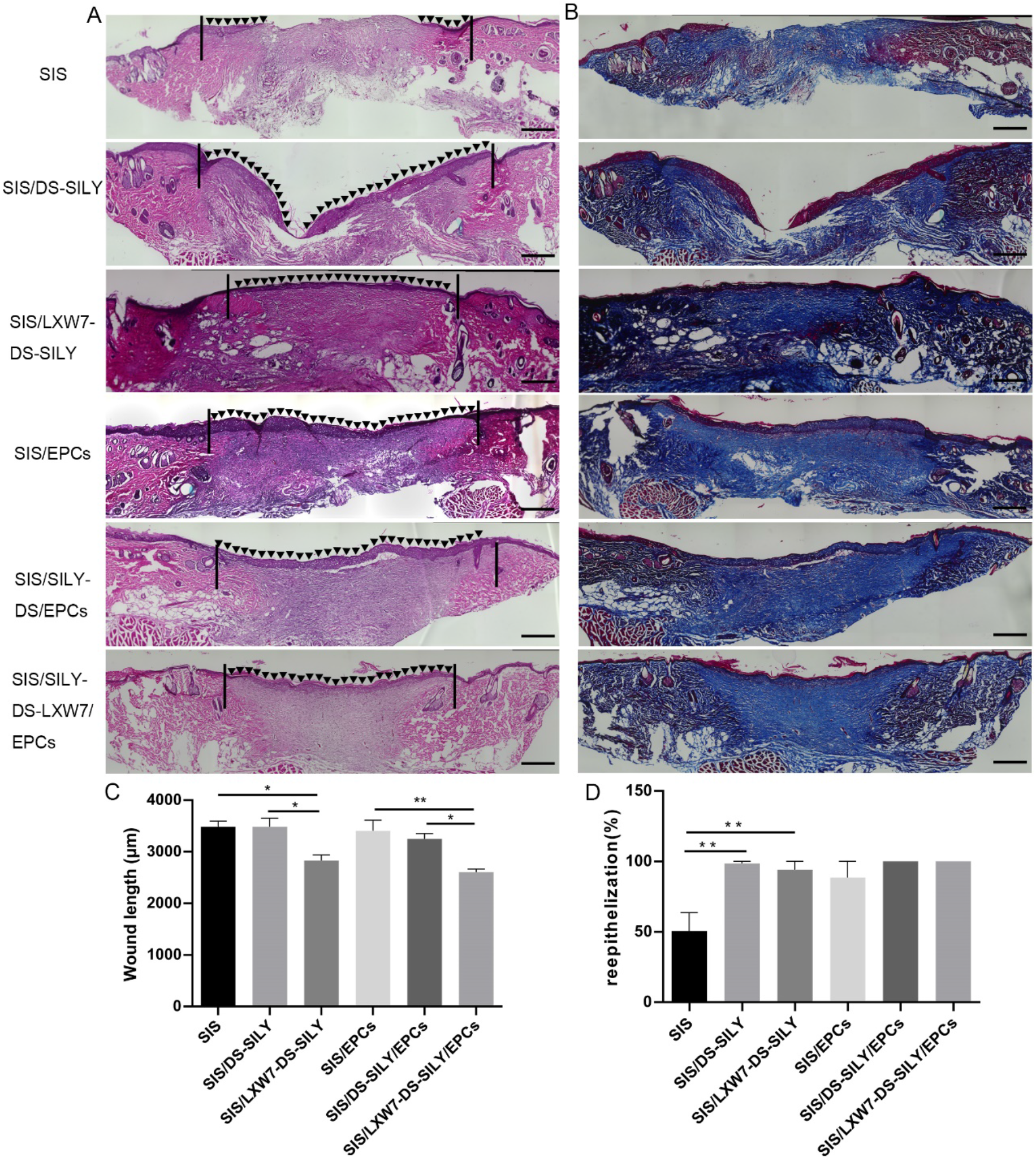
**A**. Representative images of H&E staining of wounds at day 14 under non-ischemic condition. **B**. Representative images of Masson Trichrome staining of wounds at day 14 **C-D**. Quantification of the wound length (C) and percentage of re-epithelization (D) of different modified SIS scaffolds. Scale bar = 500 µm. Data are expressed as mean ± SEM. **p<0.01, *p<0.05, n=5 per group.

#### 3.4.3 LXW7-DS-SILY modified SIS scaffolds promoted neovascularization and stimulated collagen deposition in non-ischemic wound areas

To evaluate neovascularization, we morphometrically assessed blood vessel density at wound sites using immunostaining for rat anti-endothelial cell antibody-1 (RECA-1) and alpha-smooth muscle actin (α-SMA) at day 14 (Figure 6A). Wounds treated with SIS/LXW7-DS-SILY scaffold showed a significantly higher vessel number than SIS/DS-SILY and SIS scaffolds in the regenerated tissue (Figure 6D). When wounds were treated with different scaffolds seeded with ZDF-EPCs, SIS/LXW7-DS-SILY/EPCs promoted neovascularization compared to control scaffolds with ZDF-EPCs (Figure 6D). In addition, blood vessels in the SIS/LXW7-DS-SILY/EPCs group displayed a mature structure with a larger lumen when compared to control groups (Figure 6A).

**Figure 6.**
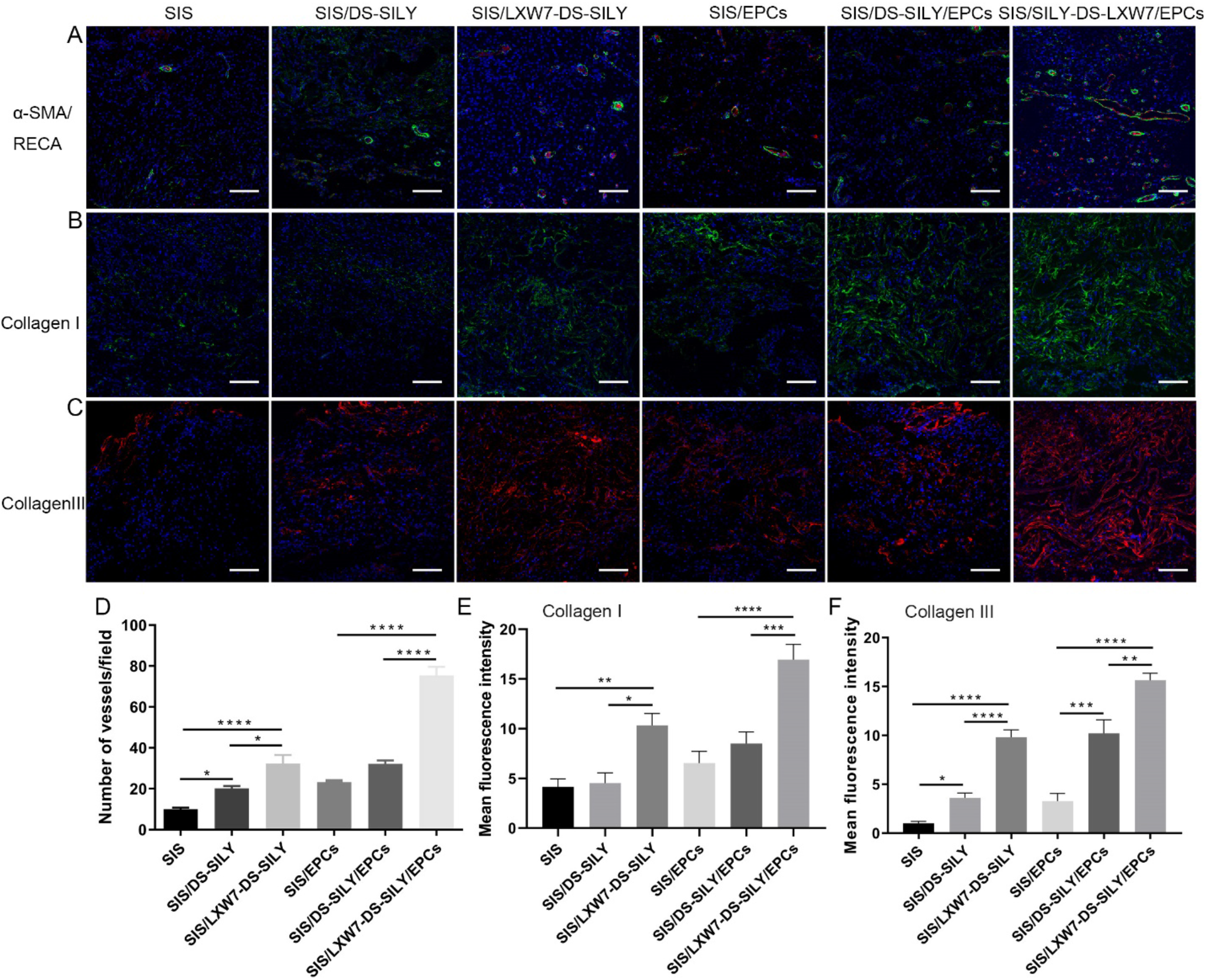
Neovascularization and collagen deposition evaluation of wounds treated by different scaffolds under non-ischemic condition. **A**. Blood vessels stained with α-SMA (green) and RECA (red) in wound bed at day 14 post surgery. **B-C**. Immunostaining of collagen I and collagen III expression in wound bed at day 14 post surgery. **D-F**. Quantification of the number of blood vessels/field (D), immunofluorescence intensity of collagen I (E) and collagen III (F). Data are expressed as mean ± SEM. **p<0.01, *p<0.05, n=3 per group.

To observe collagen morphology and distribution, Masson Trichrome staining was conducted. Masson staining showed arrangement of newly formed collagen in the regenerated tissue in all six groups (Figure 5B). On day 14, densely packed and basket-weave patterns typical of collagen fibers were observed for the SIS/LXW7-DS-SILY/EPCs group while only sparse, loosely packed collagen fibers were observed in the SIS-only group (Figure 5B)

To detect whether the collagen deposition could be stimulated by the various treatment groups, collagen immunostaining of the tissue sections was performed. Enhanced collagen I and collagen III staining intensity was observed in SIS/LXW7-DS-SILY/EPCs treated wounds on day 14, compared to the other wounds (Figure 6B, 6C). SIS/LXW7-DS-SILY also showed a significantly higher collagen I and III staining intensity compared to SIS/DS-SILY and SIS groups (Figure 6E, 6F).

#### 3.4.4 LXW7-DS-SILY modified SIS scaffolds accelerated diabetic ischemic wound healing and supported ZDF-EPC survival

To detect the effect of the different scaffolds on wound healing under ischemic conditions, we treated the ischemic wounds with SIS/LXW7-DS-SILY/EPCs, SIS/DS-SILY/EPCs, SIS/EPCs, SIS/LXW7-DS-SILY, SIS/DS-SILY or SIS. Digital photographs were obtained 0, 3, 7, 11, and 14 days after the operation (Figure 7A). The wound area in all groups decreased over time and the average observed wound closure rate was significantly higher in the SIS/LXW7-DS-SILY group (44.1±0.05%), compared with SIS group (30.0±0.06%) at day 14 (Figure 7B panel a). There was no difference between the SIS/DS-SILY (46.3±0.08%) and the SIS/LXW7-DS-SILY group (44.1±0.05%) or SIS group (30.0±0.06%). With respect to all groups loaded with ZDF-EPCs, the wounds treated with SIS/LXW7-DS-SILY/EPCs showed a wound closure rate of 61.6±0.05%, significantly higher than that of the SIS/DS-SILY/EPCs (43.5±0.05%) and SIS/EPCs (34.8±0.05%) groups. There was no difference between SIS/DS-SILY/EPCs (43.5±0.05%) and SIS/EPCs groups (34.8±0.05%) (Figure 7B panel b). Finally, the SIS/LXW7-DS-SILY/EPCs group showed a significantly higher healing rate than the SIS/LXW7-DS-SILY group highlighting the importance of the ZDF-EPCs in ischemic wound healing (Figure 7B panel c).

**Figure 7.**
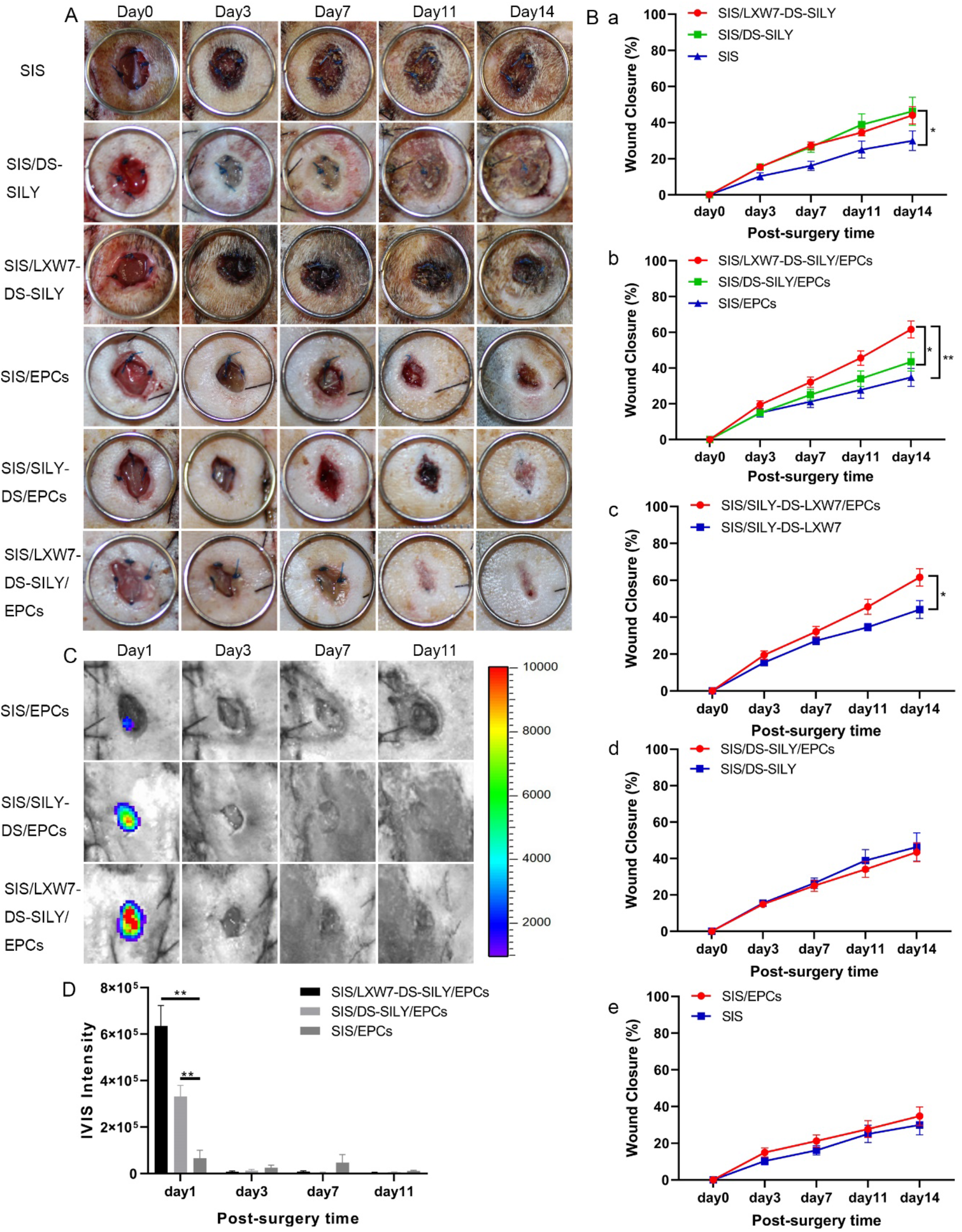
SIS/LXW7-DS-SILY constructs accelerate wound closure and support survival of ZDF-EPCs under ischemic condition. A. Representative images of healing in wounds treated with different groups at day 0, 3, 7, 11 and 14. **B**. Quantification of the wound closure rates of different groups. Data are expressed as mean ± SEM. SIS, SIS/LXW7-DS-SILY, SIS/EPCs group: n=8; SIS/DS-SILY: n=7; SIS/DS-SILY/EPCs: n=11; SIS/LXW7-DS-SILY/EPCs: n=12. C. Bioluminescence image of wounds transplanted with different modified scaffolds loaded with ZDF-EPCs. D. Quantification of the bioluminescence signals intensity (Most of the SIS scaffolds spontaneously fall off the wounds by day 11). Data are expressed as mean ± SEM. n=5 per group. **p<0.01, *p<0.05.

To investigate cell survival of transplanted cells in ischemic wounds, Td-Tomato/luciferin-labeled ZDF-EPCs were seeded on different scaffolds and imaged by IVIS (Figure 7C). The bioluminescence signal in SIS/LXW7-DS-SILY/EPCs group only showed a significantly higher intensity when compared to SIS/EPCs group at day 1. The intensity in all three groups decreased to baseline by day 11 (Figure 7D).

#### 3.4.5 LXW7-DS-SILY modified SIS scaffolds accelerated wound healing and re-epithelialization in ischemic wound areas

Overall, the wound healing quality of ischemic wounds is much worse than in non-ischemic wounds (Figure 8A). The SIS/LXW7-DS-SILY/EPCs group showed a trend toward more rapid re-epithelialization as compared to the SIS/EPCs group, as determined by shorter residual wound length (Figure 8C, 8D). However, no significant differences were found between the SIS/LXW7-DS-SILY/EPCs and SIS/DS-SILY/EPCs groups. Without EPCs, SIS/LXW7-DS-SILY scaffolds alone still have a positive effect on re-epithelialization and wound length when compared to the SIS group.

**Figure 8.**
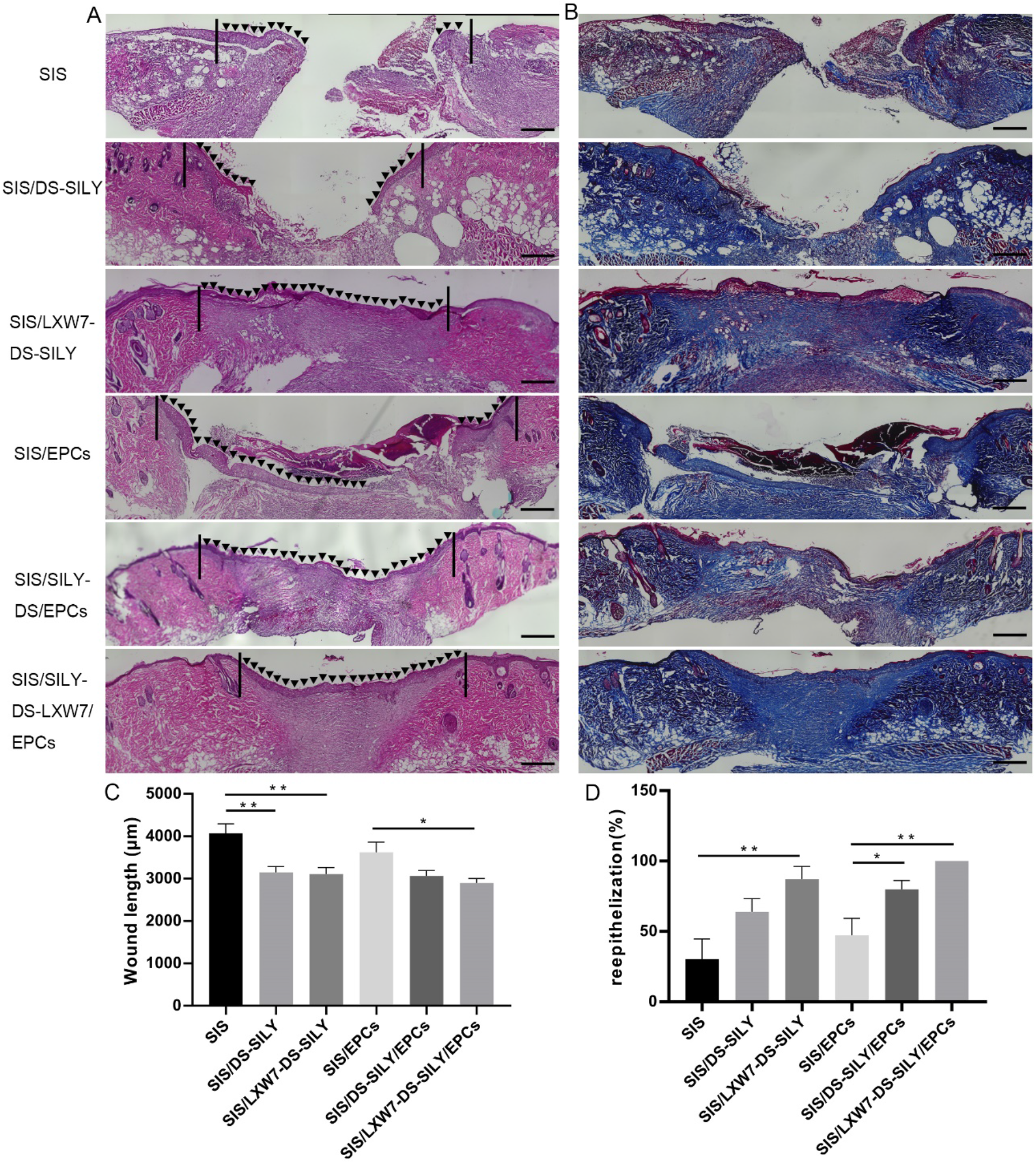
**A**. Representative images of H&E staining of wounds at day 14 under ischemic condition. **B**. Representative images of Masson Trichrome staining of wounds at day 14 **C-D**. Quantification of the wound length (C) and re-epithelization rates (D) of different groups. Scale bar = 500 µm. Data are expressed as mean ± SEM. **p<0.01, *p<0.05, n=5 per group.

#### 3.4.6 LXW7-DS-SILY modified SIS scaffolds promote angiogenesis and stimulate collagen deposition in ischemic wounds

To evaluate ischemic wound angiogenesis, we stained tissue sections for RECA-1 and α-SMA to assess capillary densities at day 14 (Figure 9A). Wounds treated with SIS/LXW7-DS-SILY scaffolds showed a significantly higher vessel number than SIS/DS-SILY and SIS scaffolds in the regenerated tissue (Figure 9D). When wounds were treated with different scaffolds seeded with ZDF-EPCs, SIS/LXW7-DS-SILY/EPCs promoted angiogenesis more than the other two groups (Figure 9D).

**Figure 9.**
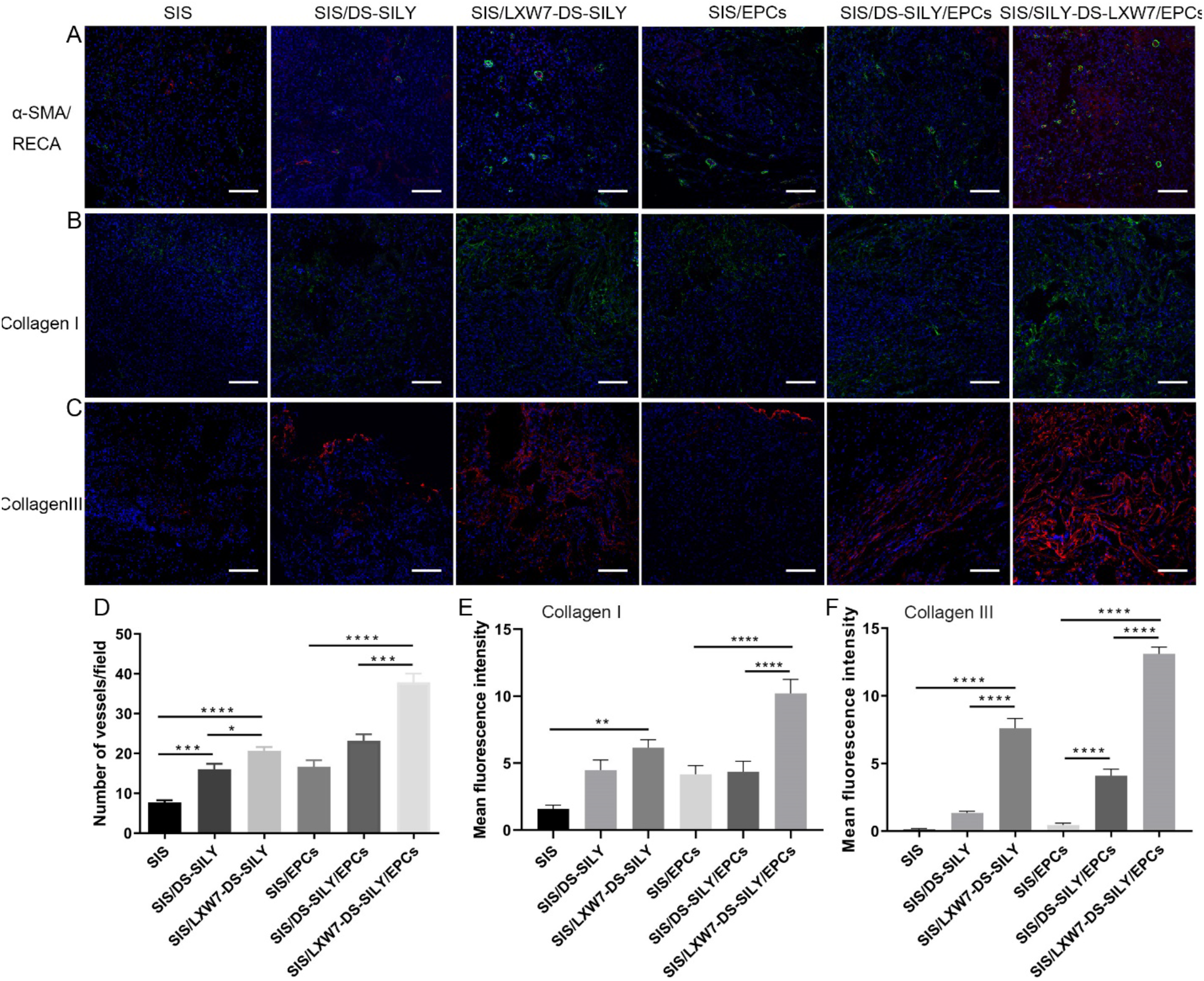
Neovascularization and collagen deposition evaluation of wounds treated by different scaffolds under ischemic condition. **A**. Blood vessels stained with α-SMA (green) and RECA (red) in wound bed at day 14 post surgery. **B-C**. Immunostaining of collagen I and collagen III expression in wound bed at day 14 post surgery. **D-F**. Quantification of the blood vessel number/field (D) and immunofluorescence intensity of collagen I (E) and collagen III (F). Data are expressed as mean ± SEM. ****p<0.0001, ***p<0.001, *p<0.05, n=3 per group.

The result showed that irregular and fewer collagen fibers were formed during this time point for the SIS group as compared to other groups (Figure 8B). Enhanced collagen I and collagen III staining intensity was observed in SIS/LXW7-DS-SILY/EPCs treated wounds on day 14 when compared to the other wounds (Figure 9B, 9C). SIS/LXW7-DS-SILY showed a significantly higher collagen III intensity than the SIS/DS-SILY and SIS groups, and a higher collagen I intensity than the SIS group (Figure 9E, 9F).

## 4 Discussion

It is well established that impaired healing of diabetic wounds is affected by several factors, such as hypoxia, dysfunction of fibroblasts and epidermal cells, impaired angiogenesis and neovascularization, and high levels of metalloproteases [25]. Hyperinsulinemia in diabetic patients impairs the proliferation and tube formation of EPCs [26]. Stem cells and regenerative materials exhibit potentially therapeutic effects towards wound and tissue regeneration. EPCs play an especially important role in the wound healing process by promoting angiogenesis and facilitating wound closure [5]. Hence, finding an effective way to augment the function of EPCs isolated from diabetic patients and delivering them to the wound site will be critical for accelerating the healing of diabetic wounds via autologous therapy.

The streptozotocin (STZ)-induced diabetic rodent model with non-ketosis hyperglycemia is the most widely used model for diabetic wound healing research [27]. However, the STZ model represents type 1 diabetes and has limitations such as organ toxicity, low stability, and impaired weight gain in animals [28]. Since type 2 diabetes account for the majority of patients with diabetes, and given the prevalence of ischemic wounds in type 2 diabetic patients, animal models that better recapitulate the pathophysiological situations of type 2 diabetic ischemic wounds serve as a better option for evaluating the function of innovative treatments for diabetic ischemic wound healing. In this study, we designed and developed a regenerative bioscaffold (SIS/LXW7-DS-SILY) loaded with EPCs and tested the therapeutic efficacy by localized topical application of the scaffold at excisional wound sites that were created to mimic the highly ischemic condition of the clinical DFU. We chose to use the Zucker diabetic fat (ZDF) rat model to evaluate the therapeutic effects of the regenerative treatment for diabetic ischemic wounds. The ZDF rat is a well-characterized model of the metabolic syndrome and the pathophysiologic mechanism of this model is attributed to the missense mutation of leptin receptor [28, 29]. ZDF rats exhibit obesity with diabetes and are characterized by hyperglycemia, insulin resistance, hyperlipidemia and impaired glucose tolerance [28], accompanied with delayed wound healing, impeded blood flow, and diabetic nephropathy [30-32]. ZDF rats become obese at around 4 weeks of age and develop type 2 diabetes at 8-12 weeks of age [28]. Additionally, in order to mimic the highly ischemic condition of the clinical DFU in this diabetic rat model, we inserted a silicone sheet beneath the pedicle flap containing the excisional wound to ensure no reperfusion from underlying tissue [23].

The wound healing mechanisms between rodents and humans are different due to anatomical variations. In rodents, healing occurs by wound contraction because of the paniculosus carnosus layer that is situated directly below the skin. In contrast, wound healing in humans occurs via re-epithelization and granulation tissue formation [34]. However, Slavkovsky, et al. [31] showed that the contraction of diabetic wounds in ZDF rats is impaired, and wound healing predominantly occurs by re-epithelialization at the wound site. To mimic this in our study, scaffolds were immobilized to the wounds with silk threads which can reduce wound contraction or enlargement. Together, the ZDF rat model developed in this study is an effective model to study the therapeutic efficacy of functionalized scaffolds for human diabetic ischemic wound healing.

Using this model, we showed that under the non-ischemic condition, the SIS/LXW7-DS-SILY/EPCs group displayed a significantly faster rate of wound closure and shorter wound lengths but showed no difference in the re-epithelialization process compared to SIS/DS-SILY/EPCs and the SIS/LXW7-DS-SILY groups. Under the ischemic condition, SIS/LXW7-DS-SILY/EPCs group again displayed a significantly faster rate of wound closure and shorter wound lengths, and in addition a rapid re-epithelialization compared to other groups. These results together suggest that LXW7 and EPCs improve the wound healing process. From our previous studies, we found that the LXW7-modified small diameter vascular grafts significantly promoted EPC/EC recruitment and rapidly achieved endothelialization [24]. Our results obtained here suggest that LXW7 may aid in the recruitment of endogenous EPCs in the non-ischemic condition and increases the EPC survival in the functionalized scaffold. It is also possible that the EPCs secrete paracrine factors that stimulate proximal ECs, and thus promote angiogenesis and improve wound healing in both conditions [35]. Interestingly, the healing of the SIS/LXW7-DS-SILY/EPCs group is significantly better than the SIS/LXW7-DS-SILY group under the ischemic condition but shows no difference under the non-ischemic condition. SIS/LXW7-DS-SILY alone facilitated improved healing effect under the non-ischemic condition. Because we know that LXW7 can effectively recruit endogenous EPCs [24], it is reasonable to speculate that LXW7 can promote wound healing by recruiting endogenous EPCs to wound sites under sufficient blood supply. In highly ischemic diabetic wound sites, there is insufficient blood supply; therefore, endogenous EPCs cannot be recruited as effectively to the wound. Transplanted EPCs may also serve to help in ischemic wound healing via releasing factors that support angiogenesis of the existing blood vessels. Even though the EPCs in the SIS/LXW7-DS-SILY/EPCs group were present in high numbers up to 24 hours and decreased thereafter, the increase in healing rate suggest that their presence for even a short period of time is sufficient to initiate tissue granulation, leading to re-epithelialization. The bioluminescence intensity of all the groups decreased to baseline at day 11. This is due in part to most of the SIS scaffolds spontaneously falling off at day 11.

It is well known that wound neovascularization is essential in order to generate sufficient blood supply to the wound area for constant supply of oxygen and nutrition, the two important factors that promote healing [27]. The results obtained from the ischemic and non-ischemic wounds treated with the SIS/LXW7-DS-SILY/EPCs scaffolds showed an enhancement in angiogenesis when compared with other groups. Our previous study indicated that LXW7 can bind to EPCs and ECs via αvβ3 integrin and enhance EC biological functions by increasing phosphorylation of VEGF receptor 2 (VEGF-R2) and activating mitogen-activated protein kinase (MAPK) ERK1/2 [18]. Our *in vitro* data also showed that LXW7 and LXW7-DS-SILY modified surfaces promoted attachment and growth of ZDF-EPCs (Figure 3). The results that SIS/LXW7-DS-SILY scaffolds alone had a positive effect on angiogenesis suggest that the LXW7 ligand potentially recruited and interacted with endogenous EPCs/ECs, and thus successfully promoted angiogenesis, neovascularization and wound healing in the diabetic wounds.

Proper collagen deposition and remodeling improve skin elasticity and toughness and greatly influence the outcomes of wound healing [36]. From immunohistological analyses (Figure 9), we observed that all the modified scaffolds displayed abundant and relatively well-organized collagen fibers. Since in our previous studies, DS-SILY showed the ability to improve collagen organization [14], we expect that the modification with functional ligands DS-SILY or LXW7-DS-SILY may exert positive effects on the ECM molecules in the wound areas, such as limiting collagen degradation and maintaining the architecture of collagen fibers.

Collagen I and III are central components of the dermal ECM with collagen I constituting 80-85%, and collagen III constituting 8-11% of the dermal ECM, and both playing a vital role in wound healing [37]. Previous studies revealed that sufficient amounts of collagen III in the early wound healing process would accelerate healing and result in scarless skin. [38]. It is known that ischemic wounds induce excessive production and over-activity of MMPs in the chronic wound environment, causing rapid degradation of new granulation tissue and countering the healing process [38-40]. DS-SILY was previously shown to improve collagen organization via inhibition of MMP-mediated collagen degradation [14]. In this study, the DS-SILY and LXW7-DS-SILY ligand modified groups displayed a significantly higher intensity of collagen I or III deposition when compared to the control group, suggesting that the ligand may protect the matrix from MMP degradation, and thus suppress scar formation and mimic the wound healing process as in normal skin. Future studies will be needed to further assess this aspect of the wound healing induced with LXW7-DS-SILY

In summary, this study demonstrates a novel therapeutic approach to accelerate diabetic ischemic wound healing by application of a functionalized combinatorial scaffold consisting of three functional components: 1) LXW7-DS-SILY, a bifunctional ligand; 2) SIS-ECM scaffold; and 3) EPCs. Further studies are necessary to explore matrix deposition at different time points, to investigate the cell populations recruited by the functionalized scaffolds, and to elucidate the potential mechanisms of LXW7 supporting the survival of transplanted EPCs, EPC paracrine signaling, and explore the potential application of this therapeutic approach to other diseases and conditions. Experiments involving large animals such as pigs which are more similar to human’s skin structure and function to further validate the safety and efficacy of this therapeutic approach will be warranted before any clinical use.

## 5 Conclusions

This study has successfully demonstrated the importance of LXW7-DS-SILY functionalized SIS scaffolds in enhancing wound healing in a type 2 diabetic rat ischemic skin flap model. EPCs cultured on SIS/LXW7-DS-SILY scaffolds showed enhanced attachment and growth. In addition, the data indicated that wound closure, re-epithelialization, angiogenesis and collagen deposition were enhanced in the diabetic ischemic wounds treated with the SIS-ECM scaffold functionalized with LXW7-DS-SILY and EPCs. Thus, LXW7-DS-SILY combined with an ECM scaffold and EPCs could be a promising novel treatment to accelerate healing of diabetic ischemic wounds, thereby reducing limb amputation and mortality rates of diabetic patients.

## 6 Acknowledgements

We thank Cook Biotech Inc. for their generous gifting of SIS-ECM material. This work was partially supported by California Institute for Regenerative Medicine grant number DISC1-10516-0, and Shriners Hospitals for Children developmental research award grant number 87200-NCA-19. This work utilized the Combinatorial Chemistry Shared Resource at the UC Davis Comprehensive Cancer Center which is supported by NCI P30CA093373 Cancer Center Support Grant.

## Data Availability Statement

All data generated or analyzed during this study are included in this published article.

---

**Figure S1.**
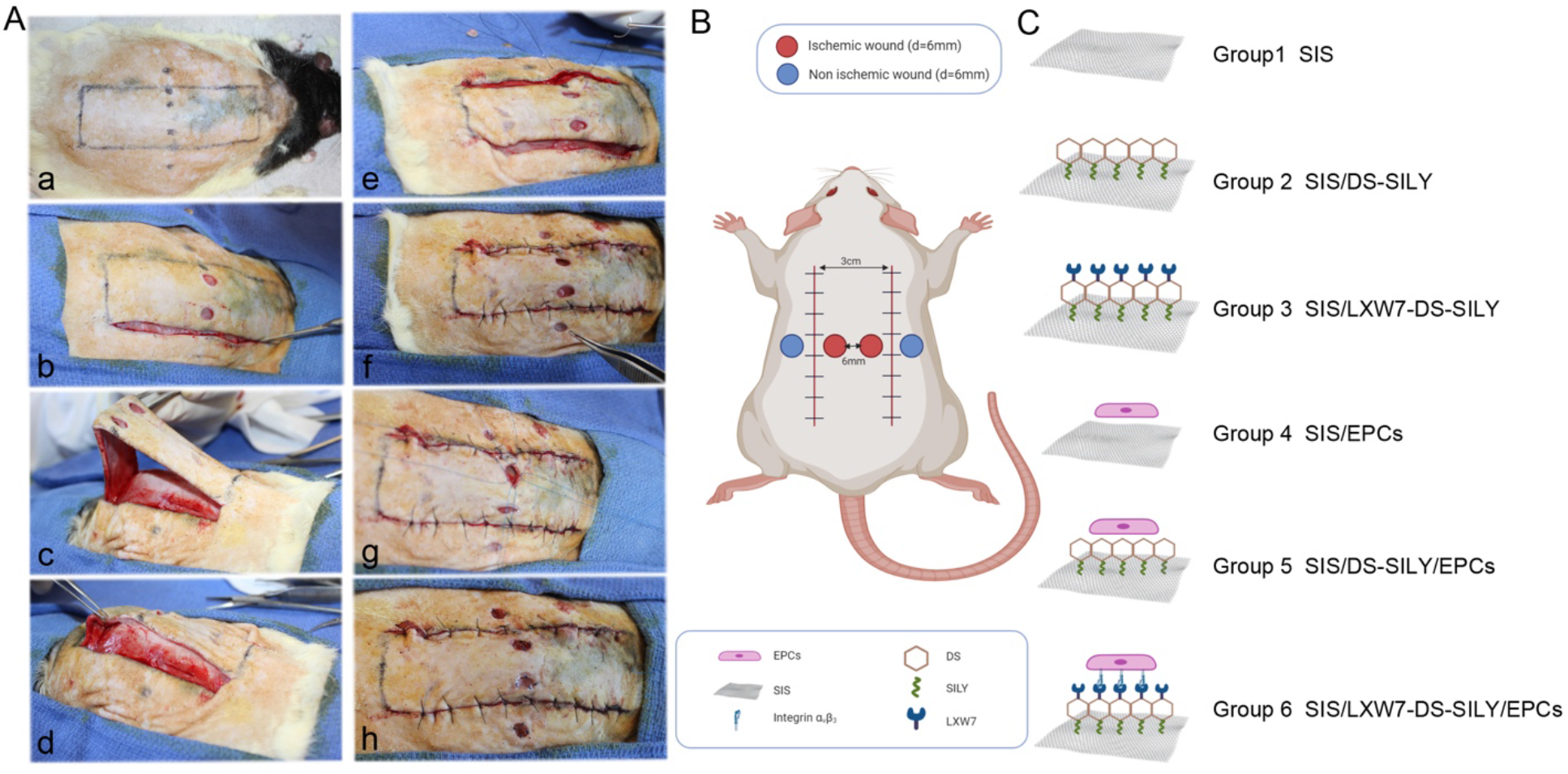
Ischemic skin flap model of Zucker Diabetic Fatty rat and experimental design. **A-B**. Demonstrates the process of making Zucker Diabetic fatty rat ischemic skin wounds model. **C**. Demonstrates the different ligand modified SIS groups used to treat the wounds.

**Figure S2.**
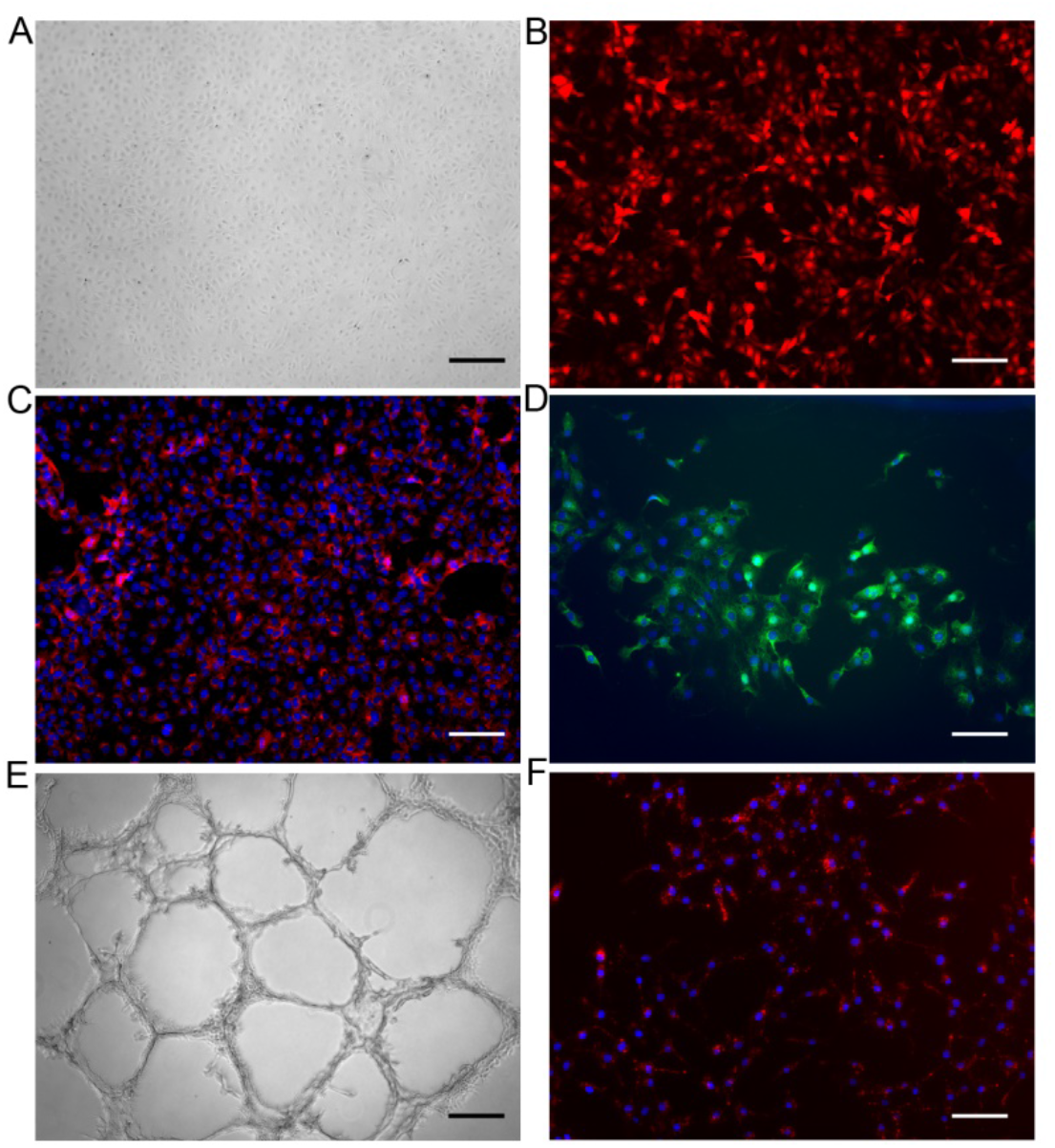
Characterization of ZDF-EPCs. A-B. Representative phase contrast. (A) and fluorescent (B) images of pCCLc-MNDU3-LUC-PGK-Tomato-WPRE lentiviral vector transduced ZDF-EPCs. **C-D**. Immunofluorescence staining results of expression of CD31 (C) and VE-Cadherin (D). **E**. *In vitro* tube formation results of ZDF-EPCs. **F**. Acetylated low-density lipoprotein uptake by ZDF-EPCs. Scale bar = 100µm (C, D, F). Scale bar = 200 µm (A, B, E).

**Figure S3.**
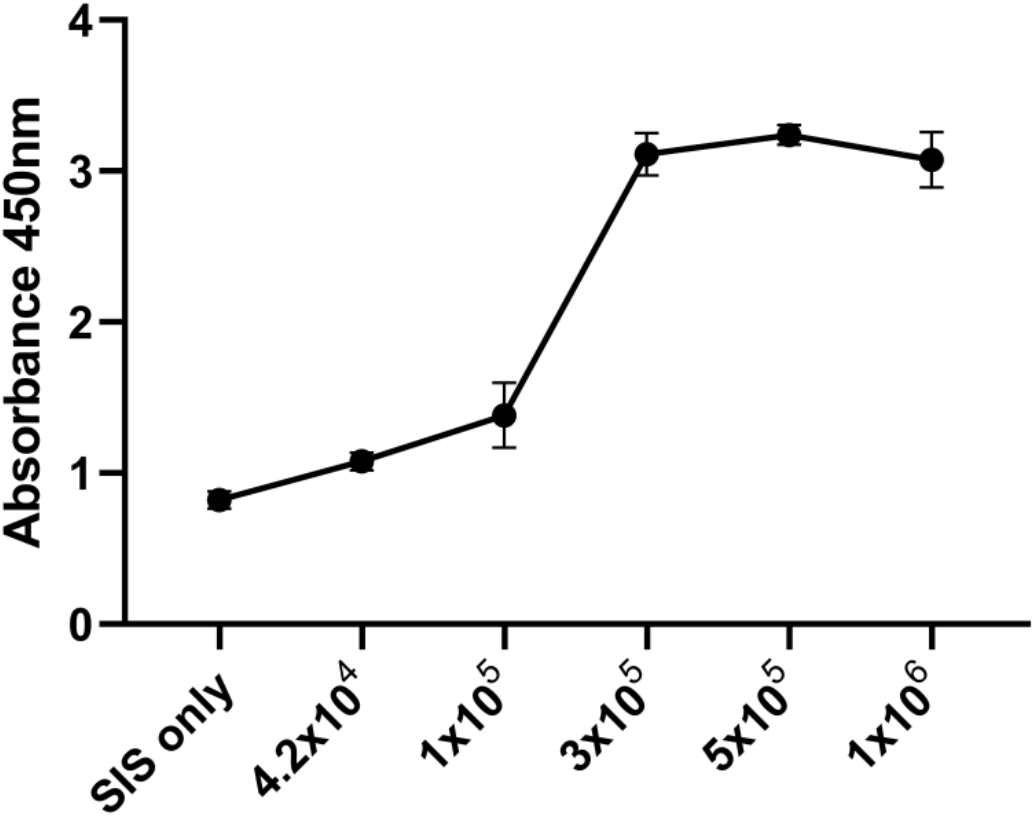
ZDF-EPCs loading density on SIS-ECM. Mean results of CCK-8 loading assay demonstrates the absorbance values of increasing concentrations of ZDF-EPCs seeded on SIS-ECM up to 5 × 10^5^ cells/cm^2^. n = 3.

## Notes

### Competing Interest Statement

KL, AP and AW are founders in VasoBio Inc, which has a license to the LXW7 peptide.

